# Sedimentation and soil carbon accumulation in degraded mangrove forests of North Sumatra, Indonesia

**DOI:** 10.1101/325191

**Authors:** Daniel Murdiyarso, Bayu Budi Hanggara, Ali Arman Lubis

## Abstract

Mangrove ecosystems are often referred to as “land builders” because of their ability to trap sediments transported from the uplands as well as from the oceans. The sedimentation process in mangrove areas is influenced by hydro-geomorphic settings that represent the tidal range and coastal geological formation. We estimated the sedimentation rate in North Sumatran mangrove forests using the ^210^Pb radionuclide technique, also known as the constant rate supply method, and found that mudflats, fringes, and interior mangroves accreted 4.3 ± 0.2 mm yr^−1^, 5.6 ± 0.3 mm yr^1^, and 3.7 ± 0.2 mm yr^−1^, respectively. Depending on the subsurface changes, these rates could potentially keep pace with global sea level rise of 2.6−3.2 mm yr^−1^, except the interior mangrove they would also be able to cope with regional sea-level rise of 4.2 ± 0.4 mm yr^−1^. The mean soil carbon accumulation rates in the mudflats, fringes, and interior areas were 40.1 ± 6.9 g C m^−2^yr^−1^, 50.1 ± 8.8 g C m^−2^yr^−1^, and 47.7 ± 12.5 g C m^−2^yr^−1^, respectively, much lower than the published global average of 226 ± 39 g C m^−2^yr^−1^. We also found that based on the excess of radioactive elements derived from atomic bomb fallout, the sediment in the mudflat area was deposited since over 28 years ago, and is much younger than the sediment deposited in the interior and fringe areas that are 43 years 54 years old, respectively.

## Introduction

Mangrove ecosystems provide numerous invaluable services including supporting (nutrient cycling, net primary production, and land formation), provisioning (food, fuel, and fiber), and regulating (climate, flood, storm surges, and pollution) services [1–5]. Situated in a transition zone between terrestrial and oceanic environments, mangrove forests play particularly important roles in moderating fresh water flow from the upland, while buffering against tidal ranges of the sea and saline water [6]. The unique shape and form of the root systems of mangrove species enable them to trap and accumulate sediments, which often contain large quantities of organic carbon [7–9]. This ability is influenced by local hydrology, geography, and topography [9,10]. In some cases, sedimentation is followed by colonization, expansion, and migration of pioneering mangrove species [11]. Therefore, carbon sequestration and storage above and below ground in mangrove ecosystems are effective in mitigating climate change [2,4].

The sustainability of the services that mangroves, including those in North Sumatra, provide is facing increasing pressure from aquaculture and agricultural development [12]. In addition, climate change and its impacts on sea-level rise have increased coastal vulnerability to erosion and inundation. Sea level has risen more than 5 cm over the last 20 years or 3.2 mm yr^−1^on average—a rate that has nearly doubled since 1990 [13,14]—and is expected to continue to increase in the future [15]. Although mangrove tree species are able to tolerate inundation by tides, they can die and their habitat formation can be damaged if, as a result of sea-level rise, the frequency and duration of the inundation exceeds their specific physiological thresholds [16,17]. Sedimentation in coastal areas that is faciltated by mangrove forests demonstrates that mangroves are “land builders” [7] and can adapt to rising sea levels. The rate of sedimentation can be an important factor to determine the sustainability of mangrove management.

The ^210^Pb radionuclide dating technique is employed to estimate sedimentation using a geochronology approach. ^210^Pb dating methods are an invaluable tool for recent (∼100 years) geochemical studies [18,19]. The natural ^210^Pb radionuclide is measured from each sediment interval as the ratio between ^209^Po and ^210^Pb, both radionuclides originate from the decay of uranium (^238^U) [20, 21]. In the process of decaying ^210^Pb, it is known that there are supported ^210^Pb and unsupported ^210^Pb. Supported ^210^Pb is formed by the decay of ^226^Ra in eroded parent rock and accumulates in the sediments. The level of supported ^210^Pb in an ecosystem is generally differ very little because it comes from the same parent rock. Unsupported ^210^Pb accumulates in sediments that enter the ecosystem [21]. This technique has frequently been used to investigate study short-term sedimentation and carbon accumulation rates (14,18,22,23]; however, it is not appropiate for long-term sediment accumulation (> 100 years) [24].

Rapid development in coastal areas due to antrophogenic activity is threatening vertical accretion and horizontal expansion of sediment in mangrove forests. These impacts include altered hydrological patterns, sedimentation, and nutrient loads that result from disturbances that restrict hydrological connections [14,25,26]. It is therefore important to understand the process of sedimentation, especially in degraded and threatened mangroves.

This study was designed to quantify the rates of sedimentation and carbon accumulation in degraded mangrove forests in Deli Serdang regency, a low-lying coastal zone in North Sumatra, Indonesia. The region is influenced by the effluent of the large city of Medan and the busy harbor port of Belawan (see Fig 1). Surrounded by shrimp ponds and newly developed oil palm plantations, the remaining mangroves in Sei Percut experience tremendous environmental pressure.

**Fig 1.**
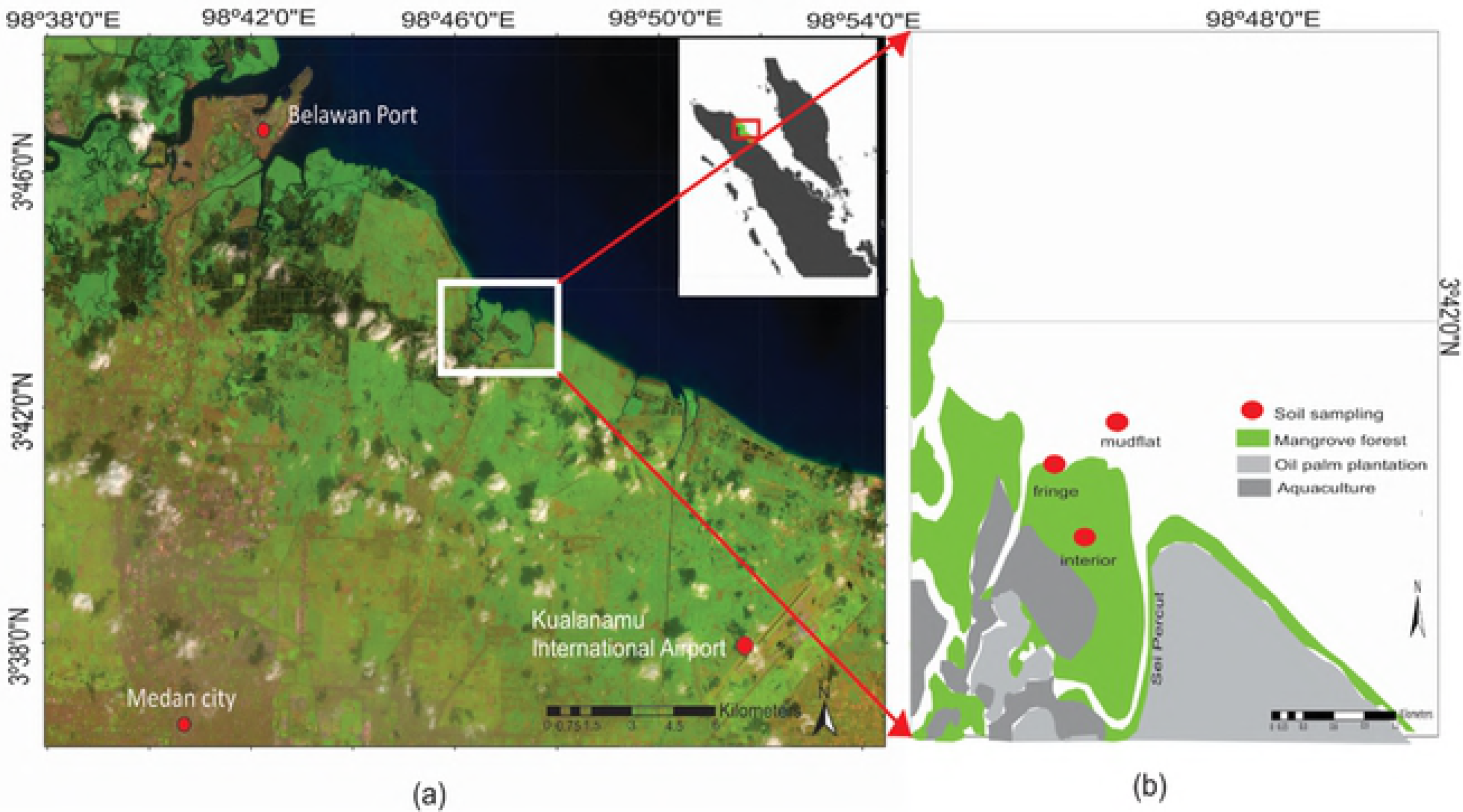
Study area of mangrove forests in Deli Serdang regency, North Sumatra, Indonesia.

The mangroves are dominated by *Avicennia* sp. and *Rhizophora* sp. Besides natural colonization of *Avicennia* sp. in the mudflat area, restoration has also been attempted, in the mudflats and in the abandoned and active ponds in the interior.

## Methods

### Site selection

The site selected is located at 3°46′15.56–3°42′53.32 N and 98°42′28.23–98°47′22.33b E. It is characterized by a monsoonal tropical climate with a mean temperature of 30°C and annual rainfall of 1848 mm [27]. The sea tide ranges from 0.9 to 2.6 m with a diurnal pattern influenced by the Andaman Sea. We selected three hydro-geomorphic settings representing mudflat, fringing mangrove and interior mangrove areas as shown in Fig 1. Three core soil samples were collected from these settings to capture the sedimentation processes and carbon storage.

### Sediment sampling and sample preparation

The sampling point in the mudflat was located approximately 15 m from the coastline (or fringe zone) and the interior at around 375 m from the coastline. Soil cores at each hydro-geomorphic setting were collected to a depth of 50 cm and sliced at 2 cm intervals for the first 10 cm and then 5 cm intervals (see Fig 2).

**Fig 2.**
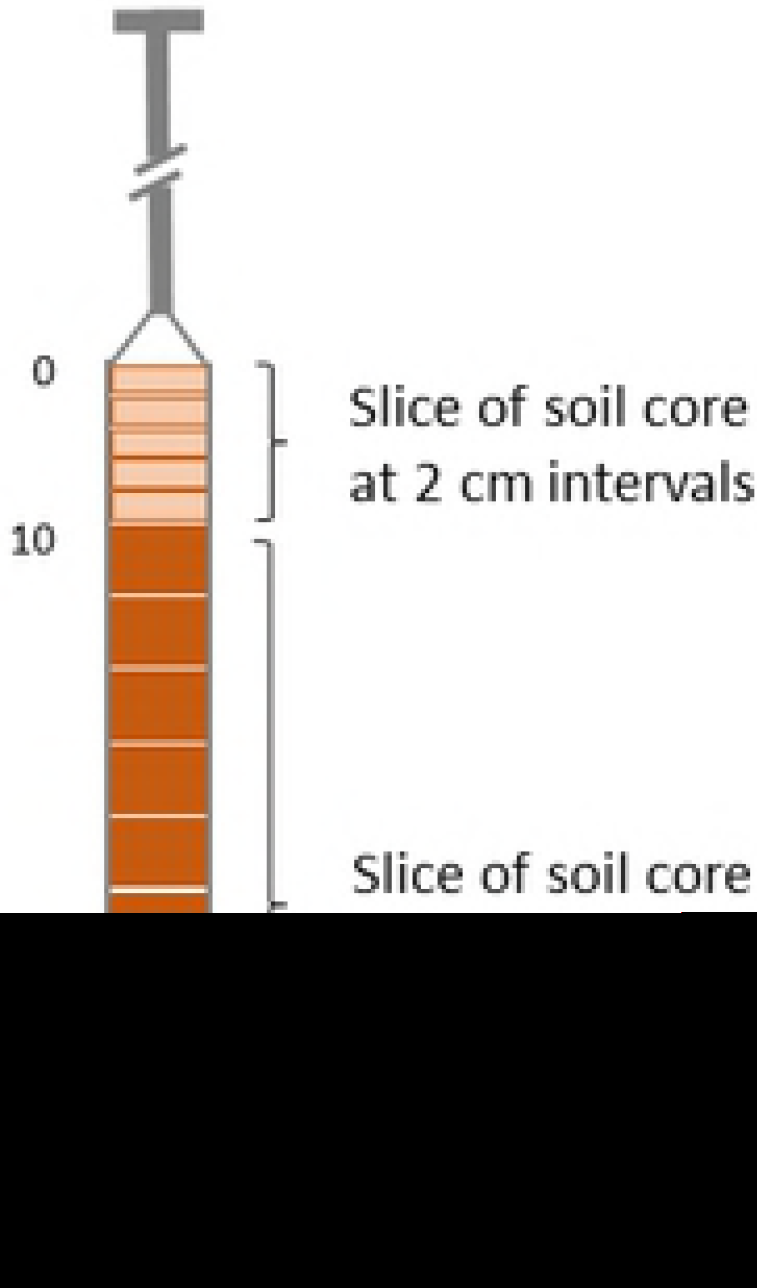
Soil core for sediment sampling in the first 50 cm below the surface.

The samples were oven-dried at a temperature of 40°C (to avoid oxidation of carbon) until a constant weight was reached. Each sample was homogenized using a mortar and pestle. Samples were dried, crushed and sieved through 120 (ϕ = 0.125 mm) mesh size (cohesive sediment). Roots and other material containing calcareous sediments were removed. Bulk density was determined for each interval by dividing the dry weight by the sample volume. The analysis of carbon content of the sediments was carried out employing a combustion method using *TruSpec Analysis CHNS* for each layer.

### Sediment accumulation rate

The sediment accumulation rates were calculated using a constant rate of supply (CRS) method [21]. Approximately 5 g dried sediment combined with 0.2 ml of radioactive tracer of ^209^Po were dissolved in a 10 ml of HCl (1:1), 10 ml of HNO_3_, 15 ml of H_2_O and 5-6 drops of 30% H_2_O_2_ and dried over a water bath at 80°C. A volume of 10 ml of HCl (1:1) and 40 ml of distilled water was added to the dry residue, before reheating over the water bath for approximately 10 minutes. The solution was filtered through filter paper No. 42 to remove sediment and rinsed with 30 ml of 0.3N HCl. The filtrate was then dried over the water bath. Volumes of 4 ml of HCl (1:1) and 50 ml of 0.3N HCl were added to the dry residue. Then 400 mg of ascorbic acid was added to complex out any dissolved iron present that might interfere with the plating processing of the Po isotopes. The sediment solution was then plated onto a 2.2 cm diameter copper disk. Po isotopes were quantified using a Canberra Alpha Spectrometer, Model 7401 with Passivated Implanted Planar Silicon detector Type A450 20AM. Measurements were carried out until a Gaussian spectrum was obtained (standard deviation was less than 10%).

To account for variability in sedimentation throughout time, the CRS model, developed by Appleby [21], was employed to calculate sediment age.

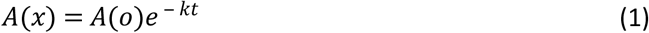

where *A(x)* is the unsupported ^210^Pb activity below the individual segment being dated (Bq kg^−1^), *A(o)* is the total unsupported ^210^Pb activity in the soil column (Bq kg^−1^), *k* is the ^210^Pb decay constant (0.0311 yr^−1^), and *t* is the age of sediments (yr) at each segment. This can be obtained from:

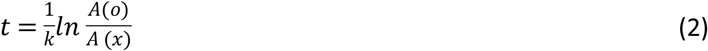

The constant supply of unsupported ^210^Pb (Bq m^−2^), *C*, was used to estimate sediment accumulation rate, *r* (kg m^−2^yr^−1^) at a certain segment and calculated as:

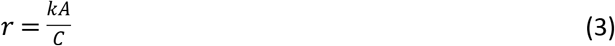

To obtain the sediment accretion rate (mm yr^−1^), the sediment accumulation (g m^−2^ yr^−1^) was divided by soil the bulk density, *BD* (g cm^−3^).

Following Marchio et al. [22], the soil carbon accumulation rate, C_acc_ (g C m^−2^ yr^−1^) was calculated for each hydro-geomorphic setting as:

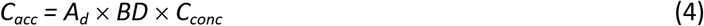

Where *A*_*d*_ is the sediment accretion rate at a certain layer (cm yr^−1^), and *C*_*conc*_ (g C g-soil^−1^) is the average soil carbon concentrations of the same layer.

## Results

### Bulk density and soil carbon content

Bulk density (*BD*) and soil carbon content are shown in Fig 3. Their variation with depth in all hydro-geomorphic locations showed opposing trends, with *BD* increasing with depth, while carbon content decreased. *BD* ranged between 0.36 g cm^−3^ and 0.77 g cm^−3^ with the widest range found in the interior (0.36-0.74 g cm^−3^), followed by the mudflat (0.43-0.77 g cm^−3^). The narrowest range was found in the fringing mangrove (0.40-0.55 g cm^−3^). In general, the variation of *BD* is within the range of most mangrove forests found across Indonesia [4].

**Fig 3.**
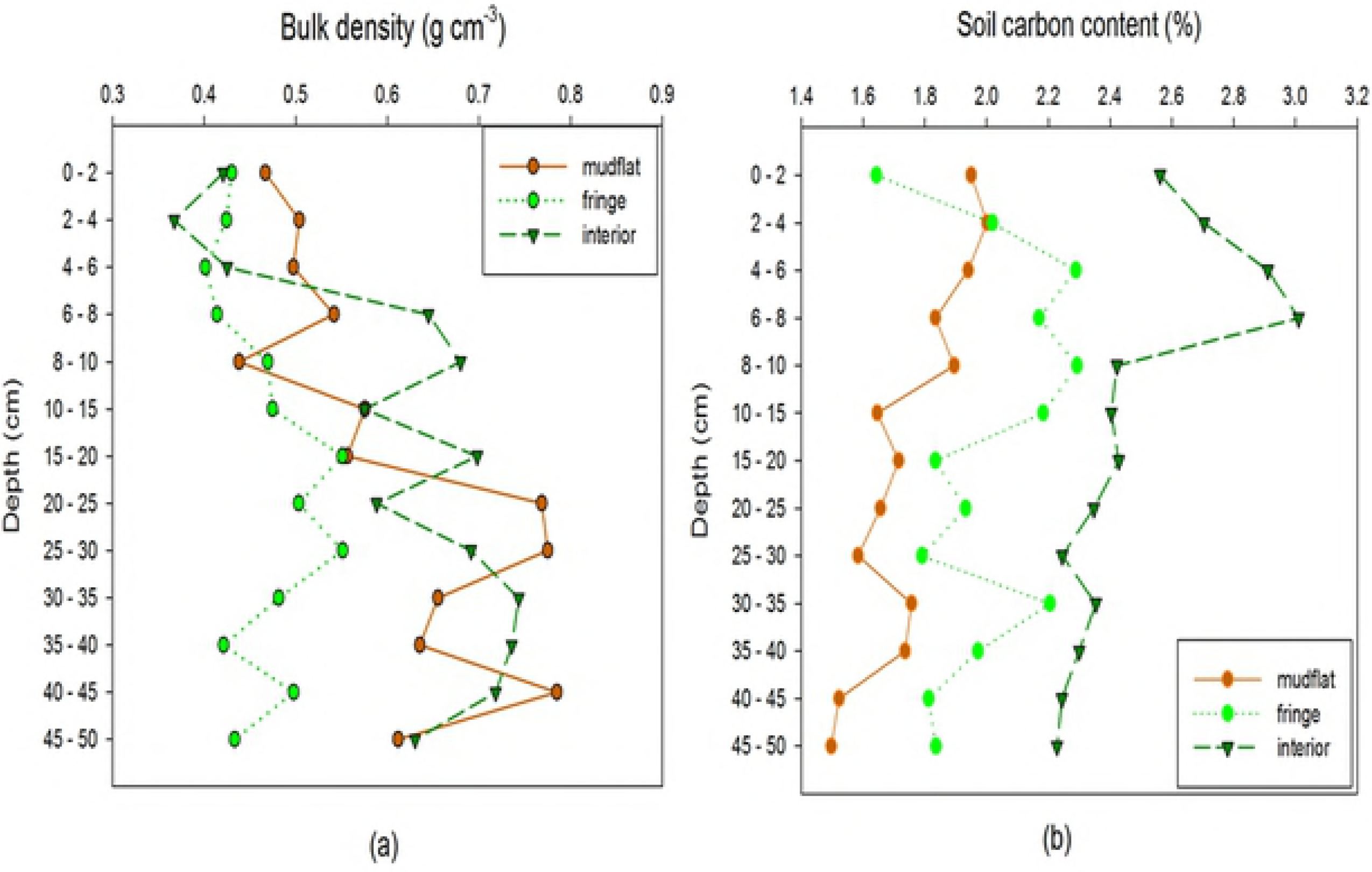
Bulk density (g cm^−3^) and soil carbon content (%C) at different hydro-geomorphic settings in mangrove forests in Deli Serdang, North Sumatra.

On average, mudflat and interior mangroves had similar *BD*s of 0.60 ± 0.11 g cm^−3^ and 0.60 ± 0.12 g cm ^−3^, respectively, with much lower *BD* in the fringing mangrove of 0.46 ± 0.05 g cm^−3^. These values suggest that the hydro-geomorphic setting dictates how sediments settle, are compacted and disturbed (or protected) by the hydrodynamics of the sea and fluvial water.

The average soil carbon content of Deli Serdang mangroves across the hydro-geomorphic setting ranged between 1.5% (mudflat) and 3.0% (interior), with fringe mangroves having an intermediate carbon content of 2.0%. This distribution signifies the role of mangrove vegetation as the primary source of carbon. Deposited and decomposed litter and decayed fine roots contribute to the *in-situ* organic carbon. These figures are far lower than the figures found in undisturbed mangroves in Sumatra (9.4%), Kalimantan (9.7%), and Sulawesi (15.6%). Moreover, they are even lower than degraded mangroves on Java of 5.6% [4].

### ^210^Pb radionuclide activity and sediment age

The unsupported ^210^Pb activity in Deli Serdang mangroves is shown in Fig 4. The mudflat area values ranged between 44.0 ± 1.8 Bq kg^−1^ and 66.9 ± 3.7 Bq kg^−1^. In the fringe mangroves, they ranged between 35.1 ± 1.8 Bq kg^−1^ and 75.7 ± 4.2 Bq kg^−1^, while the interior mangroves ranged from 39.9 ± 2.2 Bq kg^−1^ to 78.7 ± 4.3 Bq kg^−1^.

**Fig 4.**
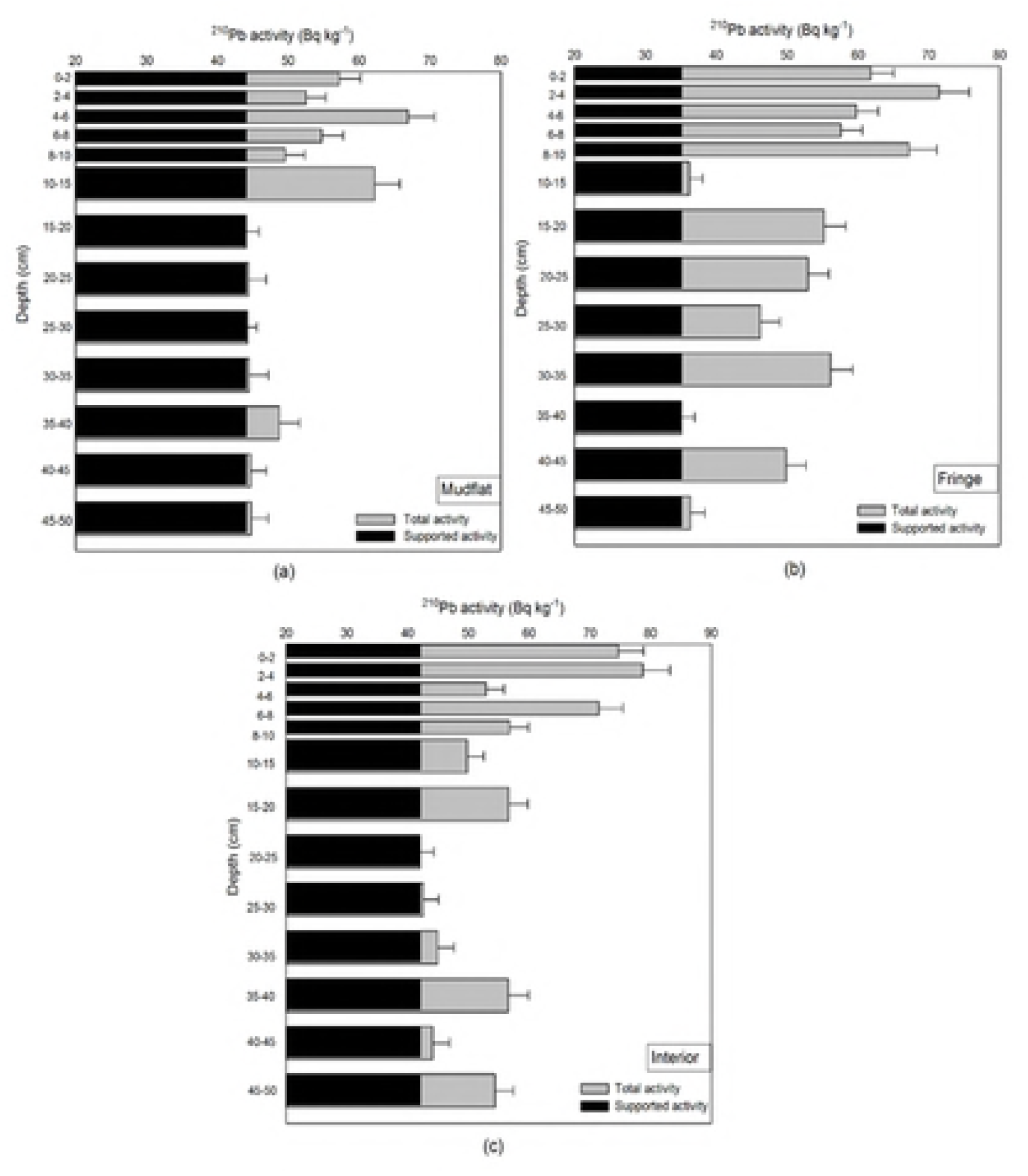
^210^Pb activity in the sediment across hydro-geomorphic setting of mangrove forests in Deli Serdang, North Sumatra: (a) mudflat, (b) fringe, (c) interior. The grey bars represent total activity of ^210^Pb (supported + unsupported), whereas the black bars represent supported ^210^Pb activivty derived from the lowest activity found in the sample. The supported ^210^Pb was formed from decayed ^226^Ra in eroded parent rock and accumulated in the sediments, and the unsupported ^210^Pb was formed from decayed ^222^Rn in nature and accumulated in the sediment.

The total ^210^Pb activity in the mudflat and fringe in the upper layer fluctuated because there was no vegetation binding the sediment on the mudflat area, so they are subject to the influence of currents and waves producing fluctuations in the suspended sediment in the area. In the fringe area, the variation was due to the influence of watersheds and less well-established root systems of colonizing mangrove seedlings.

The supported ^210^Pb activity values were obtained from the lowest unsupported values [28]. Their variation across hydro-geomorphic settings was relatively small. The values in the mudflat, fringe, and interior were 44.0 ± 3.7 Bq kg^−1^, 35.1 ± 2.3 Bq kg^−1^, and 42.1 ± 1.9 Bq kg^−1^, respectively. However, due to the limitations of CRS analysis the depth of the core that could be analysed was up to 8 cm, 25 cm and 10 cm for the mudflat, fringe and interior settings. Nevertheless, the CSR is the least arguable method compared with mass balance method, as it measures the vertical decline of ^210^Pb concentration and provides a chronology of sedimentation for up to the past century [29].

The estimates of sediment age in each layer and at all hydro-geomorphic settings are shown in Fig 5. The oldest sediment was found in the fringing mangroves, which was formed 54.7 ± 3.2 yrs ago (around 1961) at 25 cm deep, followed by the interior mangroves of 43.1 ± 2.3 yrs (around 1973) at a depth of 10 cm. The youngest sediment was formed 28.4 ± 1.6 yrs ago (around 1988) found in the mudflat area at 8 cm deep. These ages of sedimentation in North Sumatra are much younger than those found in Everglades National Park in the Gulf of Mexico, where the sedimentation processes began in 1926 [23]. Since the region was very much affected by high storm surges, the sediment deposits and carbon burial were relatively higher.

**Fig 5.**
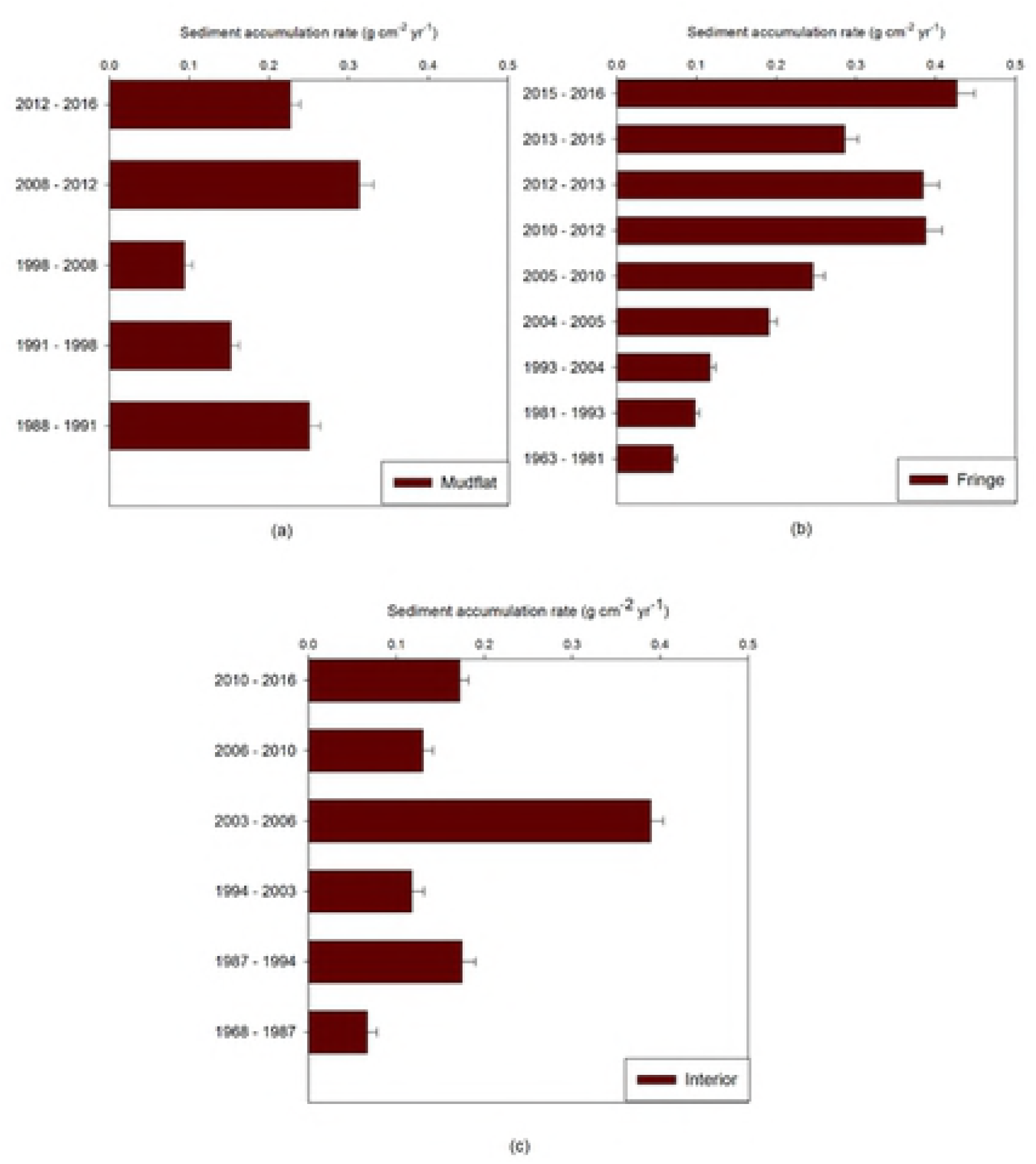
Sediment accumulation rate and the age of sediment in each hydro-geomorphic setting of mangrove forests in Deli Serdang, North Sumatra: (a) mudflat, (b) fringe, (c) interior. The oldest sediment was found in the fringe mangroves of about 55 years.

### Sediment accretion and soil carbon accumulation rates

The constant supply of unsupported ^210^Pb is used to estimate sediment accumulation rate (mass per unit area) or sediment accretion rate (thickness). This is a good indicator of growth capacity, horizontal expansion of mangrove ecosystems, and, to some extent, effectiveness as carbon sinks nd disturbance regimes [23]. Fig 6a shows sediment accretion and soil carbon accumulation rates in all hydro-gemorphic settings with fringing mangroves being the largest (5.6 ± 0.3 mm yr^−1^), followed by mudflats at 4.3 ± 0.2 mm yr^−1^, and interior mangroves at 3.7 ± 0.2 mm yr^−1^. This pattern is associated with the size, shape, and spatial distribution of trees. The rate increases with increasing density of trees especially those of *Rhizophora sp*. and their complex root systems, which not only trap the mud but most likely also allows them to withstand greater sedimentation. This was particularly the case for fringing mangroves as the interior mangroves were isolated from receiving sediments. The sedimentation in the mangroves of Deli Serdang, North Sumatra was generally due to marine and river sedimentary deposits. These results are in line with previous work, which generally found that sedimentation rate was the highest along the coastline and decreased in the area near the mainland [11].

**Fig 6.**
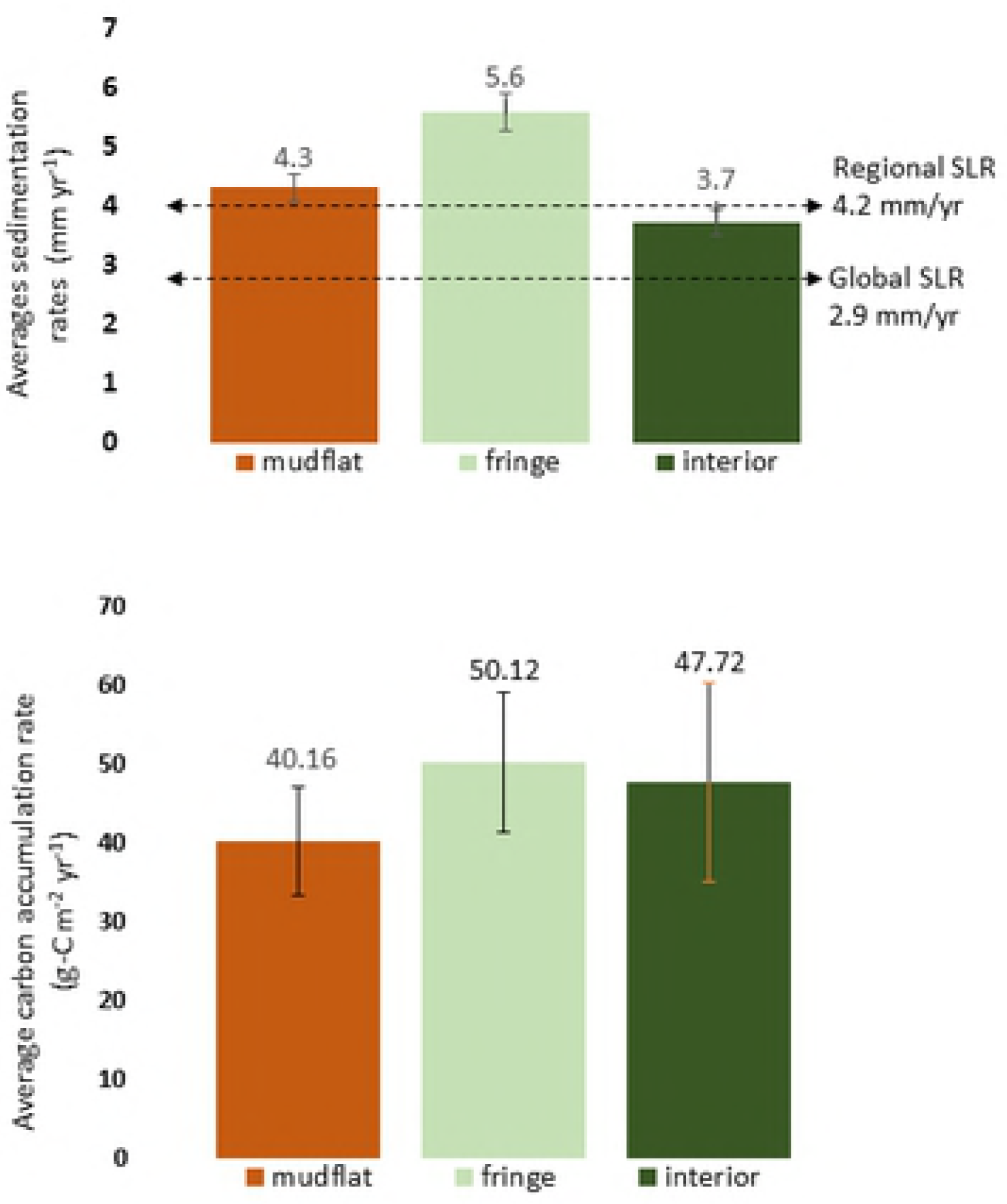
Sediment accretion rate (a) and soil carbon accumulation rate (b) in each hydro-geomorphic setting of mangrove forests in Deli Serdang, North Sumatra: mudflat, fringe, and interior. It is shown in (a) that sedimentation rate in all settings can cope with global sea level rise (SLR) but only interior mangroves cannot cope with regional SLR.

The accumulation rate of soil carbon depends on sedimentation rate, *BD*, and the carbon concentration of sediment. As shown in Fig 6b the average value of soil carbon accumulation in the mudflat area was 40.1 ± 6.9 g C m^−2^yr^−1^. In the fringe mangroves, it was 50.1± 88.4 g C m^−2^yr^−1^, and in the interior mangroves was 47.7 ± 12.5 g C m^−2^yr^−1^. Although mudflat zone demonstrates higher sedimentation rate than interior mangroves, the accumulation of carbon is lower. This is because the presence of mangrove forest is an important source of autochthonous carbon. However, the rate of carbon burial across the hydro-geomorphic shown in Deli Serdang mangroves are very low compared with the global average of around 226 ± 39 g C m^−2^yr^−1^ [30].

## Discussion

Reduction of land-based emissions, such as reducing emissions from deforestation and forest degradation, known as REDD+ is often promoted as climate mitigation measures. Mangrove forest is a unique example by which climate change mitigation can be simultaneously implemented with adaption strategies. Combining the two roles could even be more attractive for stakeholders at national level to demonstrate the nationally determined contributions (NDCs) as stipulated in the Paris Agreement, where climate change adaptation and mitigation are implemented in a balanced manner. Moreover, sub national and local agenda could even be built around adaptation strategies that usually gain less attention in both programming and budgeting processes.

### Mangroves promote adaption to rising sea level

The global average sedimentation rate in mangrove forests ranges between 0.1 and 10.0 mm yr^−1^, with a median value of 5 mm yr^−1^ [31, 32]. Based on ^210^Pb radionuclide analysis using the CRS method we found an average rate of sediment accretion rate in the mudflat area of 4.3 ± 0.2 mm yr^−1^, while in the fringe mangroves the rate was 5.6 ± 0.3 mm yr^−1^, and the lowest rate was found in the interior mangroves of 3.7 ± 0.2 mm yr^−1^. Assuming that no sub surface changes took place, sedimentation in Deli Serdang mangrove forests can keep pace with global sea-level rise of 2.6-3.2 mm yr^−1^ [33] and regional sea-level rise of 4.2 ± 0.4 mm yr^−1^ [15], except for the interior mangroves, which would not withstand the regional sea-level rise.

However, various processes can cause changes in mangrove sediment, including surface and subsurface processes. Previous studies [34, 35] describe processes occurring on or above the surface of mangrove soils, including sedimentation (deposition of material to the soil surface), accretion (binding of this material in place), and erosion (loss of surface material). We did not monitor the sub surface processes, such as growth and decomposition of roots, soil swelling and shrinking associated with moisture content, and compaction, compression, and rebound of soils due to changes in the weight of the overlying material. In addition, at a larger scale, subsidence or geological movement may affect the sediment [36]. These mean that sediment accretion alone is not the best predictor of mangrove forests′ resilience or capacity to adapt to sea-level rise. Measurement using the rod surface elevation table combined with a marker horizon may give further information [37].

Up to 80% of the sediments delivered by the tides are retained in mangrove forests [32, 38], enhancing mangrove colonization and expansion. The morphology and root systems of mangrove vegetation with their strong and complex shapes facilitates the trapping of sediment particles along the tidal ranges in the coastal area [38]. Natural regeneration, therefore, should be promoted.

### Effectiveness of mangroves in mitigating climate change

Globally, organic carbon burial in coastal wetlands, including mangrove ecosystems, varies hugely depending on carbon content of the sediment deposited, net primary production, root and microbial activities, tidal waves and hydrodynamics, and hydro-geomorphic settings. It is generally accepted that the burial rate of soil sediments in mangrove ecosystems accumulates carbon between 163 and 265 g C m^−2^yr^−1^ [22,31,39–42]. These are the largest among any terrestrial ecosystems after saltmarsh [39].

Although carbon accumulation from sediment in mangrove forests in Deli Serdang, North Sumatra was very low (40-50 g C m^−2^yr^−1^), seem to have significantly low carbon burial rates, it was reported that mangroves in Hinchinbrook Channel, Australia has an even lower average rate of 26 g C m^−2^yr^−1^ [43]. In contrast, an extremely high rate of 949 g C m^−2^yr^−1^ was found in Tamandare, Brazil [44].

The colonization, recruitment, and expansion of mangrove species vegetation in newly reclaimed land could gradually increase carbon sequestration through photosynthesis by the mangroves and litter deposition. Rehabilitation or restoration of mangrove to recover the carbon storage capacity of the ecosystem will take long time; therefore conserving intact mangroves is crucial and more effective to mitigate climate change.

These observations suggest that in terms of management, there is no one single solution for restoring degraded mangroves. A combination of methods and objectives could be explored to answer particular site-specific challenges. To some extent ecosystem services capable of accommodating a number of objectives and interests of multiple stakeholders. Ecosystem services provided by mangroves are among the candidates that attract local governments and community, including fishing community (nursery ground, coastal protection and pollution filtering), urban community (regulating micro-climate, cultural and education objects).

## Acknowledgements

This study was part of the Sustainable Wetlands Adaptation and Mitigation Program (SWAMP) funded by the United States Agency for International Development (*AID-BFS-IO-17-00005*). The field work facilitated by the Yagasu Foundation and their field staff is gratefully acknowledged. The immense support of the staff at the Center of Isotopes and Radiation Application, National Nuclear Agency is recognized and thanked. Finally, we thank two anonymous reviewers for the very helpful comments that greatly improved this manuscript.

## Author Contributions

**Conceptualization:** Daniel Murdiyarso.

**Data curation:** Daniel Murdiyarso.

**Formal analysis:** Daniel Murdiyarso and Bayu Budi Hanggara.

**Funding acquisition:** Daniel Murdiyarso.

**Investigation:** Daniel Murdiyarso and Bayu Budi Hanggara.

**Methodology:** Daniel Murdiyarso, Bayu Budi Hanggara, and Ali Arman Lubis.

**Project administration:** Daniel Murdiyarso.

**Supervision:** Daniel Murdiyarso and Ali Arman Lubis.

**Validation:** Daniel Murdiyarso and Ali Arman Lubis.

**Visualization:** Daniel Murdiyarso and Bayu Budi Hanggara.

**Writing ± original draft:** Daniel Murdiyarso, Bayu Budi Hanggara, and Ali Arman Lubis.

**Writing ± review & editing:** Daniel Murdiyarso

